# Convergent approaches to delineate the metabolic regulation of tumor invasion by hyaluronic acid biosynthesis

**DOI:** 10.1101/2022.06.08.495338

**Authors:** Adrian A. Shimpi, Matthew L. Tan, Michael Vilkhovoy, David Dai, L. Monet Roberts, Joe Kuo, Lingting Huang, Jeffrey D. Varner, Matthew Paszek, Claudia Fischbach

## Abstract

Metastasis is the leading cause of breast cancer-related deaths and often driven by invasion and cancer-stem like cells (CSCs). Both the CSC phenotype and invasion have been associated with increased hyaluronic acid (HA) production. How these independent observations are connected, and which role metabolism plays in this process remains unclear due in part to the lack of convergent approaches that integrate engineered model systems, computational tools, and cancer biology. Using microfluidic invasion models, metabolomics, computational flux balance analysis (FBA), and bioinformatic analysis of patient data we investigated the functional links between the stem-like, invasive, and metabolic phenotype of breast cancer cells as a function of HA biosynthesis. Our results suggest that CSCs are more invasive than non-CSCs and that broad metabolic changes caused by overproduction of HA play a role in this process. Accordingly, overexpression of hyaluronic acid synthases (HAS) 2 or 3 induced a metabolic phenotype that promoted breast cancer cell stemness and invasion *in vitro* and upregulated a transcriptomic signature that was predictive of increased invasion and worse survival in patients. Collectively, this study suggests that HA overproduction leads to metabolic adaptations that help satisfy the energy demands necessary for 3D invasion of breast cancer stem cells further highlighting the importance of engineered model systems and multidisciplinary approaches in cancer research.

## Introduction

Despite advancements in treatment options, breast cancer remains the second leading cause of cancer-related deaths in women,^1^. Mortality in breast cancer is driven by metastasis and relapse, during which cancer cells in the primary tumor invade into surrounding tissues and disseminate into distant sites to form secondary tumors. The pathogenesis of metastasis can be attributed to intratumoral heterogeneity, where phenotypic diversity enables a subset of cells to become invasive and resistant to traditional therapies^2,3^. In particular, the emergence of cancer cells with stem-like properties (CSCs) contributes to metastasis because of their self-renewal and invasive properties^4^. CSCs are identified by their expression of stem cell markers (e.g. NANOG, SOX2, OCT4, aldehyde dehydrogenase [ALDH]) and influenced by features of the tumor microenvironment including extracellular matrix (ECM)^5,6^. For example, CSC invasion into the surrounding stroma is controlled by ECM microarchitecture and stiffness^7,8^ including collagen fiber alignment at the tumor periphery^7,9,10^. However, the emergence and maintenance of the CSC phenotype and their consequences on 3D collagen invasion are poorly understood as studies often isolate tumor cells from the complex microenvironmental conditions that influence their behavior *in vivo*.

One key component of the tumor microenvironment influencing tumor cell phenotype and invasion is hyaluronic acid (HA). While much prior work has focused on how HA secreted by stromal cells regulates tumorigenesis^11–13^, cancer cells themselves also produce HA. In fact, overproduction of HA by tumor cells enriches for a CSC phenotype^14,15^ and correlates with worse patient prognosis^11,16^. Although several studies have delineated the specific signaling pathways by which HA modulates cell behavior^15,17^, excess production of HA also regulates tumor cells via biophysical mechanisms^18,19^. For example, biosynthesis and pericellular retention of HA as part of the glycocalyx allows tumor cells to invade and extravasate more effectively^20,21^. However, which metabolic adaptations tumor cells may use to increase HA-mediated tumor cell invasion and the role cancer cell stemness plays in this process remains unclear.

Aberrant cellular metabolism is a hallmark of cancer that has been independently tied to increased tumor cell invasion, stemness, and HA biosynthesis. Therefore, we speculated that metabolism may serve as an overarching regulator of HA-mediated tumor cell invasion. Specifically, we hypothesized that metabolic adaptations increase cancer cell stemness, which phenotypically can more readily satisfy the energetic demands of 3D invasion. Prior findings suggest that CSCs exhibit metabolic phenotypes that are distinct from non-CSCs^22–24^ and that effective tumor cell invasion through dense ECM requires metabolic adaptations including increased glucose uptake and ATP generation^25,26^. While increased glucose uptake due to aerobic glycolysis is a common feature of tumor cell metabolism, this typically results in the fermentation of glucose to lactate (known as the Warburg Effect) which produces ATP less efficiently than oxidative phosphorylation per glucose molecule. This energetically disadvantageous state must then encourage tumor cells to develop compensatory mechanisms to generate the necessary energy for 3D invasion.

Because increased HA production of CSCs is known to direct intermediate products of glycolysis into the hexosamine biosynthetic pathway (HBP)^17^ it is possible that the resulting metabolic rewiring enables their 3D invasion by providing alternative strategies for ATP production. However, identifying broad metabolic changes beyond conventional biochemical methods requires integrating computational approaches that can model the flow of metabolites through relevant large-scale metabolic networks, while simulating a desired metabolic phenotype for subsequent experimental validation. Here, we integrate engineered cell lines, microfabricated culture models, and computational approaches including flux-balance-analysis (FBA) to investigate the interconnectedness of the tumor cell phenotype, HA production, and metabolism and its influence on cancer cell invasion. We demonstrate that metabolic adaptations associated with increased HA production promote a stem-like phenotype in cancer cells, which can more readily satisfy the energy demands necessary for 3D invasion. Bioinformatic analysis of clinical data from publicly available datasets further indicated that these changes correlated with increased invasive potential and worse survival in patients. Collectively, our results suggest that HA overproduction due to metabolic reprogramming negatively influences prognosis in breast cancer patients by altering invasion, and further motivate the need for utilizing a multidisciplinary toolset to study intratumoral heterogeneity and its role in invasion.

## Results

### Cancer Stem-Like Cells Exhibit Increased Invasive Potential

To investigate differences in invasion between CSCs and their non-differentiated counterparts, we utilized the CSC reporter cell line GFP-NANOG MDA-MB-231 in which green fluorescent protein (GFP) expression is controlled by the NANOG promoter^27^ (Fig. 1a). Using fluorescence-activated cell sorting (FACS) GFP-NANOG MDA-MB-231 cells were sorted into GFP^Null^, GFP^Low^ (bottom 5% GFP expressing), and GFP^High^ (top 5% GFP expressing cells) populations to enrich for different stem-like states (Fig. 1a) whose phenotype was maintained over 5 days of culture (Supplementary Fig. 1). Moreover, the GFP^High^ cell population proliferated more slowly than the non-CSC GFP^Null^ population, consistent with a more quiescent phenotype (Fig. 1b). To study potential differences in the invasive phenotype of these different cell populations, we monitored tumor cell invasion in response to a morphogen gradient using a microfluidic collagen type I hydrogel model (Fig. 1c). In this setup, GFP^High^ NANOG MDA-MB-231 cells invaded into the 3D fibrillar collagen hydrogel region (Fig. 1d) of the device more readily than their less stem-like counterparts, validating that CSCs exhibit increased 3D invasive potential (Fig. 1e). Cell migration and invasion are energetically intensive processes requiring increased metabolic consumption^28^. Accordingly, inhibiting energy production with the glycolysis inhibitor 2-deoxyglucose (2-DG) reduced both collagen hydrogel invasion and random 2D migration of unsorted GFP-NANOG MDA-MB-231 cells (Fig. 1f,g). These results indicate that CSCs exhibit increased invasive and migratory potential in our experimental setup that was dependent on glycolytic metabolism.

**Figure 1:**
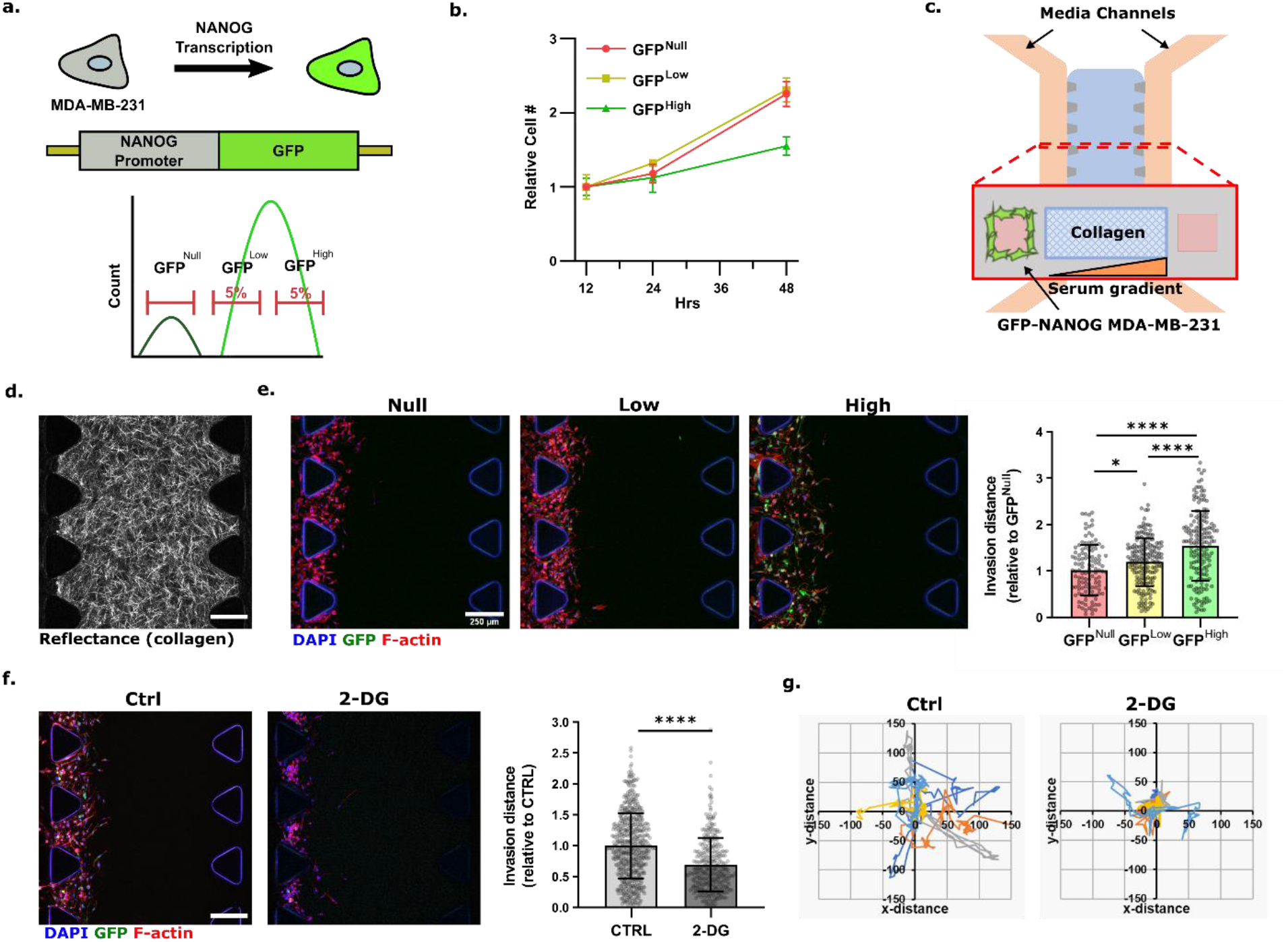
CSCs have increased invasive potential that is sensitive to metabolic challenge. **a)** Schematic of the cancer stem cell reporter line GFP-NANOG MDA-MB-231 and sorting strategy. **b)** Growth curve of sorted cells as measured by DNA amount, normalized to the first measurement at t = 12 hrs (n = 3 samples). **c)** Schematic of the microfluidic device to analyze cell invasion in response to a morphogen gradient generated by applying serum-containing medium to the right channel only. **d)** Representative confocal reflectance microscopy image of a fibrillar collagen hydrogel in microfluidic invasion device. **e)** Invasion distance of sorted GFP-NANOG MDA-MB-231 into collagen type I over 5 days (n = 4 fields of views, 1 device per condition). **f)** Invasion of GFP-NANOG MDA-MB-231 cells into collagen type I treated with or without 2-DG for 5 days (n = 3 devices per condition). **g)** Random migration of GFP-NANOG MDA-MB-231 treated with or without 2-DG for 24 hours (n = 5 representative cells for migration plots, n = 20 cells per condition for motility). Scale bar = 250 μm. * p< 0.05, ** p<0.01, *** p<0.001, **** p<0.0001

### Cancer Stem-Like Cells Have Altered Metabolism

To characterize the metabolic phenotype of cancer cells as a function of their stem-like phenotype, extracellular metabolite production and consumption rates were measured in sorted GFP-NANOG cells (Fig. 2a,b). The more stem-like cells consumed more glucose and produced more lactate relative to their non-stem-like counterparts indicative of increased glycolysis (Fig. 2a). Real-time metabolic analysis with the Agilent Seahorse (Seahorse) Analyzer of extracellular acidification rate (ECAR) and oxygen consumption rate (OCR) further suggested that CSCs not only exhibited increased glycolysis, but also oxidative phosphorylation (Fig. 2b, Supplementary Fig. 2). Acccordingly, culturing in glucose-free media decreased the percentage of GFP^High^ cells relative to media containing physiological levels of glucose (Fig. 2c). This approach also decreased the ALDH bright (ALDH^Br^) fraction (a marker for breast CSCs^29,30^) of wildtype MDA-MB-231 cells confirming that our results were not an artefact of the GFP-NANOG reporter cell line. Moreover, inhibition of glycolysis by 2-DG reduced the GFP^High^ population further corroborating that the CSC phenotype is intimately linked to glucose metabolism (Fig. 2d).

**Figure 2:**
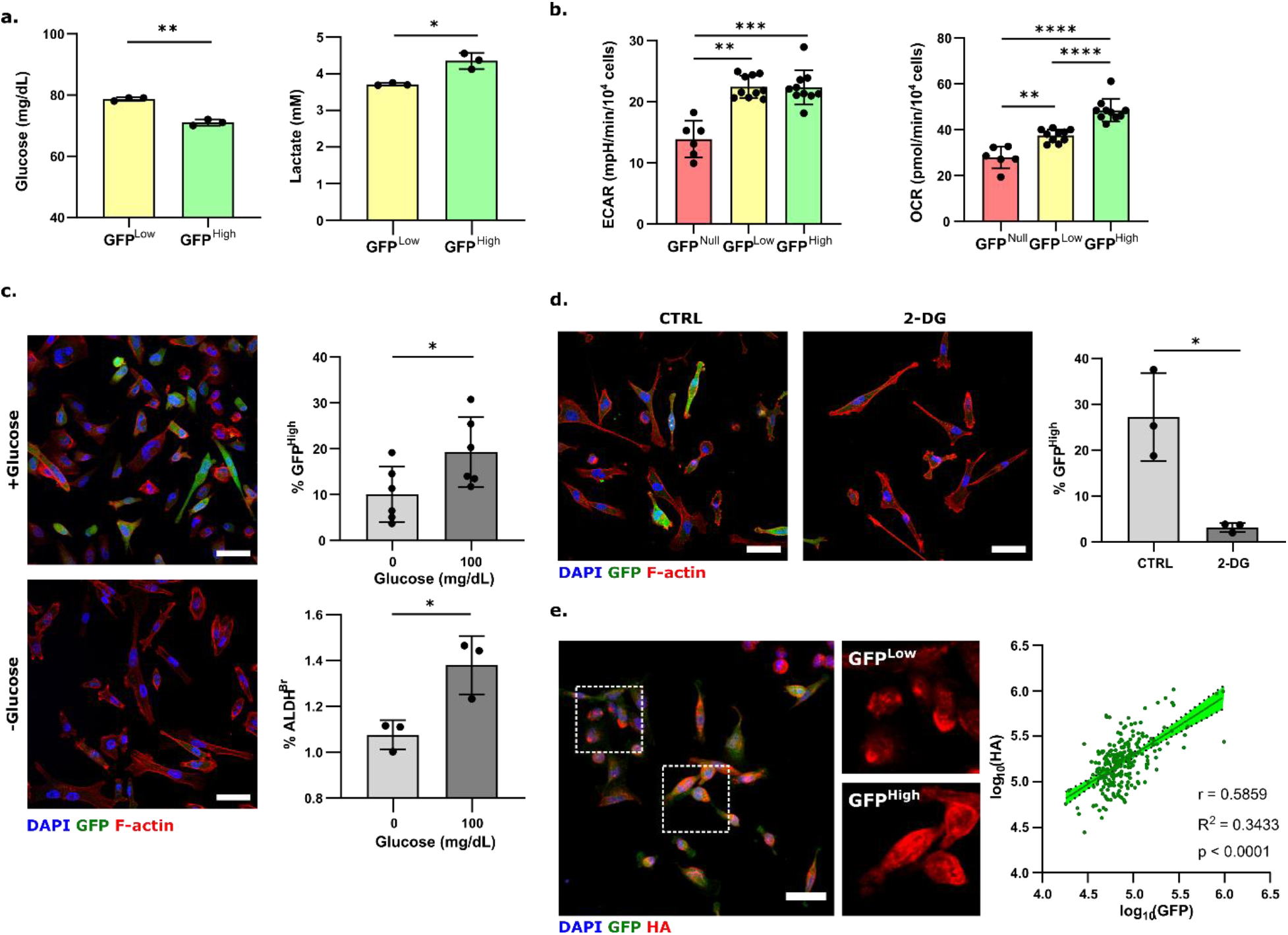
Cancer stem-like cells exhibit altered metabolism compared to non-stem-like cancer cells. **a)** Glucose and lactate concentrations in media conditioned by sorted GFP-NANOG MDA-MB-231 cells 48 hrs post sort as measured by a GlucCell glucose meter and a lactate colorimetric assay (n = 3 samples). **b)** ECAR and OCR measurements of sorted GFP-NANOG cells obtained during Seahorse-based glycolysis stress test (n > 6 samples). **c)** Fraction of GFP positive (GFP+) GFP-NANOG MDA-MB-231 (n = 6) (i) and aldehyde dehydrogenase high (ALDH+) (n = 3) (ii) parental MDA-MB-231 when cultured with 100 mg/dL glucose (+glucose) or glucose-free media (-glucose) for 72 hours. GFP+ and ALDH+ cells were determined by image analysis and Aldefluor assay, respectively. **d)** Fraction of GFP+ GFP-NANOG MDA-MB-231 cells treated with or without 2-deoxyglucose (2-DG) (n = 3). **e)** Immunofluorescence analysis of HA in the MDA-MB-231 NANOG reporter line. (n = 248 cells) Scale bar = 50 μm. * p< 0.05, ** p<0.01, *** p<0.001, **** p<0.0001.

As increased HA production has been associated with both metabolic reprogramming and the CSC phenotype^17,31^, we next assessed how HA production correlates with the CSC phenotype. Immunofluorescence (IF) image analysis identified that GFP expression in GFP-NANOG MDA-MB-231 cells positively correlated with the amount of cell surface-associated HA (Fig. 2e). These differences were likely due to changes in HA biosynthesis and subsequent cell surface retention as analysis of secreted HA by ELISA indicated low levels across all experimental conditions that were not significantly different from each other although more stem-like cells seemed to secrete slightly less HA (Supplementary Fig. 3a). Together, these results suggest that more stem-like cells exhibit increased glucose metabolism relative to their less stem-like counterparts and that these changes correlate with elevated levels of cell surface-associated HA.

### Hyaluronic Acid Production Correlates with Stemness

To better quantify the relationship between glucose metabolism, cell surface-associated HA, and stemness, GFP-NANOG MDA-MB-231 cells were stained for HA and then subjected to flow cytometry and targeted metabolomics. Consistent with the IF results, the amount of cell surface-associated HA directly correlated with GFP expression levels (Fig. 3a). Importantly, sorting the GFP-NANOG MDA-MB-231 cells into high and low HA producing cells for targeted metabolomics revealed broad changes of intracellular metabolites with hierarchical clustering separating the two HA production phenotypes (Fig. 3b). In particular, both upper glycolytic and tricarbocylic acid (TCA) cycle metabolites such as glucose-6-phosphate (G6P), fructose-6-phosphate (F6P), oxaloacetate (OAA), and citrate (CIT) were increased in the highly HA producing cells (Fig. 3c). Combining these results with the broad metabolic alterations of CSCs described above (Fig. 2), these data suggest a direct link between glycolysis-dependent HA synthesis and the CSC phenotype (Fig. 3d). To more directly test how HA synthesis affects stemness, GFP-NANOG MDA-MB-231 cells were cultured with the HA synthesis inhibitor 4-methylumbelliferone (4-MU, 0.5 mM). These results trended towards a decreased fraction of stem-like GFP^High^ cells consistent with previous studies on stemness and invasion^15,20^ (Supplementary Fig. 3b).

**Figure 3:**
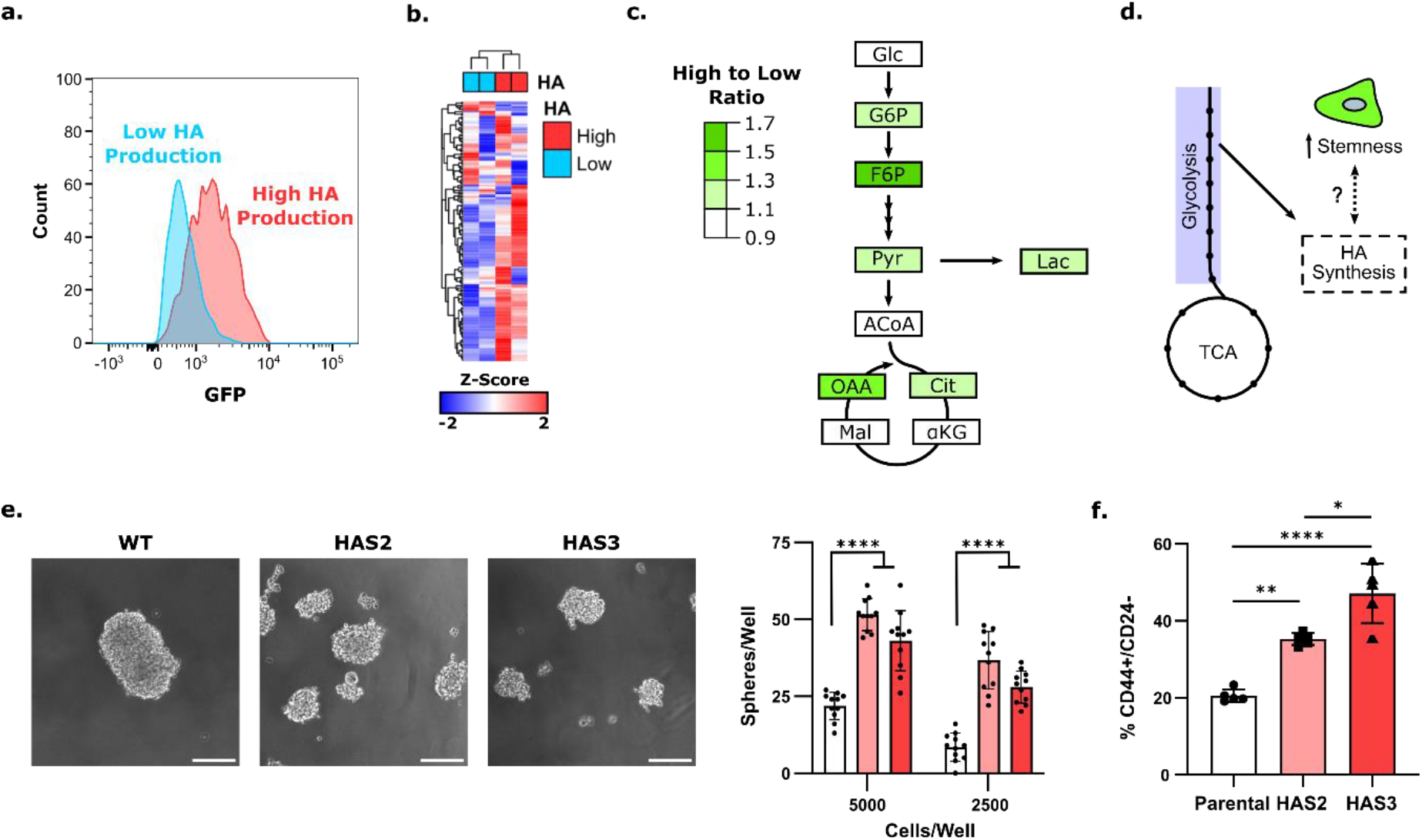
Increased HA production correlates with an increase in CSCs. **a)** Flow cytometry analysis of cellular GFP levels categorized by their levels of cell surface-associated HA. **b)** Heatmap representing changes of intracellular metabolites as measured by metabolomics of sorted HA^High^ and HA^Low^ GFP-NANOG MDA-MB-231. **c)** Graphical representation of selected metabolites in the central carbon metabolic pathway ratios between HA^High^ and HA^Low^ cells. **d)** Schematic representing the theorized relationship between glycolysis, HA synthesis, and stemness. **e)** Representative images of spheres formed through a limiting dilution assay of HAS2 and HAS3 overexpressing MDA-MB-231 and the corresponding sphere number. (n = 11) **f)** Flow cytometry analysis of the percentage of the CD44^+^/CD24^−^ population of parental or HA overproducing MCF10A cells (n = 5). * p<0.05, ** p< 0.01, **** p<0.001. Scale bar = 200 μm.

To more directly determine if increasing production of HA promotes stemness, HAS2 and HAS3 were stably overexpressed in MDA-MB-231 and the non-malignant breast epithelial cell line MCF10A. While HA is synthesized by three HAS isoforms (HAS1, 2, 3), HAS 1 varies in HA synthesis and secretion rate from HAS 2 and 3. In contrast, HAS2 and HAS3 exhibit similar sensitivities and responses to precursor availability and thus were used here^32–34^. Overexpression of HAS2/3 increased the defining characteristic of CSCs in both cell lines. More specifically, overexpression of HAS2/3 increased MDA-MB231 self-renewal as measured through sphere formation in a limited dilution assay (Fig. 3e, Supplementary Fig. 4a,b) and increased the fraction of CD44^+^/CD24^−^ MCF10A cells, which characterizes an invasive breast CSC population with more mesenchymal characteristics^35,36^ (Fig. 3f, Supplementary Fig. 4d). This population could not be assessed in HAS overexpressing MDA-MB-231 as this cell line contains an intrinsically high fraction of CD44+/CD24-cells (>85%), making changes difficult to quantify^37,38^ (Supplementary Fig. 4c). Finally, HAS3 overexpression also trended towards an increase in the ALDH^Br^ fraction of MCF10A (Supplementary Fig. 4e). Together, these results suggest that HA overproduction increases stem-like cell properties in breast cancer cells regardless of cell line and HAS2/3 isoform.

### Increased HA Production by CSCs is Associated with Increased Glucose Conversion and ATP Production

Given our results that metabolic reprogramming of CSCs supports their energy demands during invasion (Fig. 2) and that a more stem-like phenotype is associated with increased HA production (Fig. 3), we speculated that increased HA production promotes a more energetic CSC phenotype. Probing the individual contribution of HA biosynthetic pathways to the metabolic state of CSCs solely by measuring different metabolite levels, however, is challenging given the interconnectedness of most metabolic pathways. To circumvent these limitations and delineate the contribution of HA biosynthesis to CSC metabolism more directly, a flux balance analysis (FBA) model was constructed. FBA is a widely utilized mathematical approach to model the flow of metabolites (flux) through a genome-scale network of metabolic pathways, including glycolysis, oxidative phosphorylation, hexosamine biosynthesis, and amino acid consumption^39^, and has been successfully used to investigate cancer metabolism^40,41^. In contrast to traditional metabolomics, FBA also enables the simulation of a desired metabolic phenotype by adjusting model parameters such as the objective function and flux constraints. To develop the model for our study, extracellular metabolomics of GFP^High^ and GFP^Null^ cells were performed over 72 hours to define metabolite consumption/production profiles used to constrain flux values for more and less stem-like MDA-MB-231 breast cancer cells, respectively (Fig. 4a). Additionally, the objective function of the FBA model was set to maximize HA synthesis to study both the capacity of CSCs and non-CSCs to produce HA and the associated changes in metabolic flux. Results from the FBA model indicated that the more stem-like cells (GFP^High^) increased flux through the upper stages of glycolysis (i.e., conversion of glucose to fructose-6-phosphate), ATP generation, lactate production/secretion, and glutamine uptake relative to the non-stem-like cells (GFP^Null^) (Fig. 4b, Supplementary File 1). Additionally, the FBA model indicated increased flux through all HA synthesis intermediate steps for the GFP^high^ cells relative to the GFP^null^ cells (Fig. 4c). Collectively, these data suggest that the increased capacity of CSCs to synthesize HA is related to increased glucose conversion but simultaneously allows these cells to produce ATP more efficiently. To confirm the predictive value of the FBA model, a Seahorse real-time ATP rate assay was performed. This analysis verified that the mitochondrial and overall ATP production rate was increased in GFP^High^ versus GFP^Null^ cells (Fig. 4 d,e). Together, these findings suggest that the metabolic phenotype of more stem-like cancer cells leading to increased HA biosynthesis promotes ATP production by these cells. FBA models are suitable for predicting these changes.

**Figure 4:**
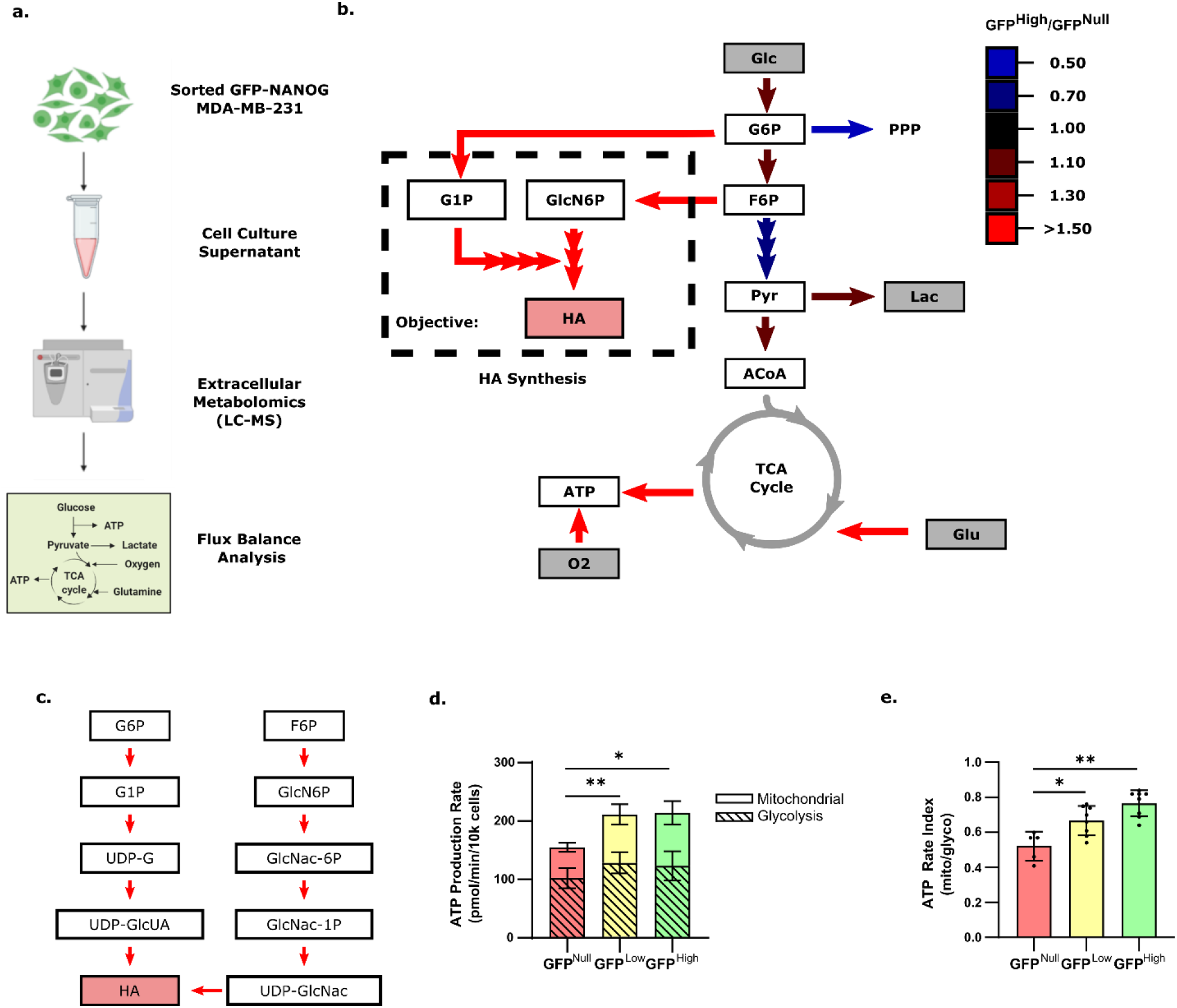
Flux balance analysis predicts increased HA production in cancer stem-like cells and ATP production. **a)** Workflow schematic. Sorted GFP-NANOG cells were cultured for 72 hrs before performing extracellular metabolomics using LC-MS. This data was used to constrain a computational flux balance analysis model. **b)** Flux balance analysis model of the bioenergetic pathway of sorted GFP-NANOG MDA-MB-231 derived from extracellular metabolomics obtained over 72 hours of culture. Fluxes shown are GFP^High^ relative to GFP^Null^. Grey metabolites indicate relevant metabolite fluxes constrained based on extracellular metabolomic profiles. Oxygen was constrained using values obtained from the Agilent Seahorse assay. For the total FBA model constraints and results, see Supplementary File 1. **c)** Expanded HA synthesis pathway flux balance analysis of GFP^High^ relative to GFP^Null^ (dashed box in B). Same legend as in B. **d)** ATP production rate and **e)**rate index derived from measurements using the Agilent Seahorse ATP real-time production rate assay kit (n ≥ 5). * p<0.05, ** p< 0.01.

### Metabolic Changes resulting from Increased HA Production Stimulate a Stem-like Breast Cancer Cell Phenotype to Promote Invasion

Our aforementioned results suggest functional links between the stem-like and invasive phenotype of tumor cells (Fig. 1), CSCs and metabolism (Fig. 2), CSCs and HA (Fig. 3), and HA and metabolism (Fig. 4). However, whether these single observations are mechanistically connected remained to be determined. Therefore, we next hypothesized that the metabolic phenotype induced by HA overproduction leads to changes in energy production that increase the stem-like phenotype of breast cancer cells to promote invasion. Indeed, Seahorse analysis revealed that MCF10A and MDA-MB-231 cells overexpressing HAS2 and HAS3 had increased ECAR compared to their parental controls suggesting an increase in glycolytic energy production (Fig. 5a). Interestingly, OCR was unchanged in HAS2/3-overexpressing MCF10A but increased in HAS2/3-overexpressing MDA-MB-231 cells (Supplementary Fig. 5a), consistent with an increase in OCR in GFP^High^ vs. GFP^Low^ and GFP^Null^ MDA-MB-231 (Fig. 2b). Treatment with 2-DG decreased both cell surface-associated and secreted HA in MCF10A cells (Supplementary Fig. 5b, c). Notably, 2-DG treatment of MCF10A cells decreased the CD44^+^/CD24^−^ fraction in the HA overproducing cells but had no effect on the parental control cells suggesting that the enhanced glycolysis associated with HA overproduction is critical to maintaining stemness (Fig. 5b). Furthermore, HA overproduction increased both total and glycolytic ATP production (Fig. 5c, Supplementary Fig. 5d) that 2-DG reversed to similar levels as in MCF10A control cells (Fig. 5c). A similar trend was noted for MDA-MB-231 cells, although 2-DG decreased glycolytic ATP production only in the HAS3 overexpressing cells. Overexpression of HAS2/3 also increased invasion of both MCF10A and MDA-MB-231 (Fig. 5d, e), consistent with their increased stem-like phenotype (Fig. 3d, e). This effect was inhibited by 2-DG treatment, implicating a functional consequence of HA-mediated stemness and metabolism in tumor cell invasion (Fig. 5d, e). As 2-DG treatment did not affect the ATP production rate in parental cells (Fig. 5c), these changes in invasion can be attributed to differences in metabolism rather than compromised cell viability. Together this data suggests that HA overproduction induces a glycolytic phenotype that is crucial for CSC-mediated invasion.

**Figure 5:**
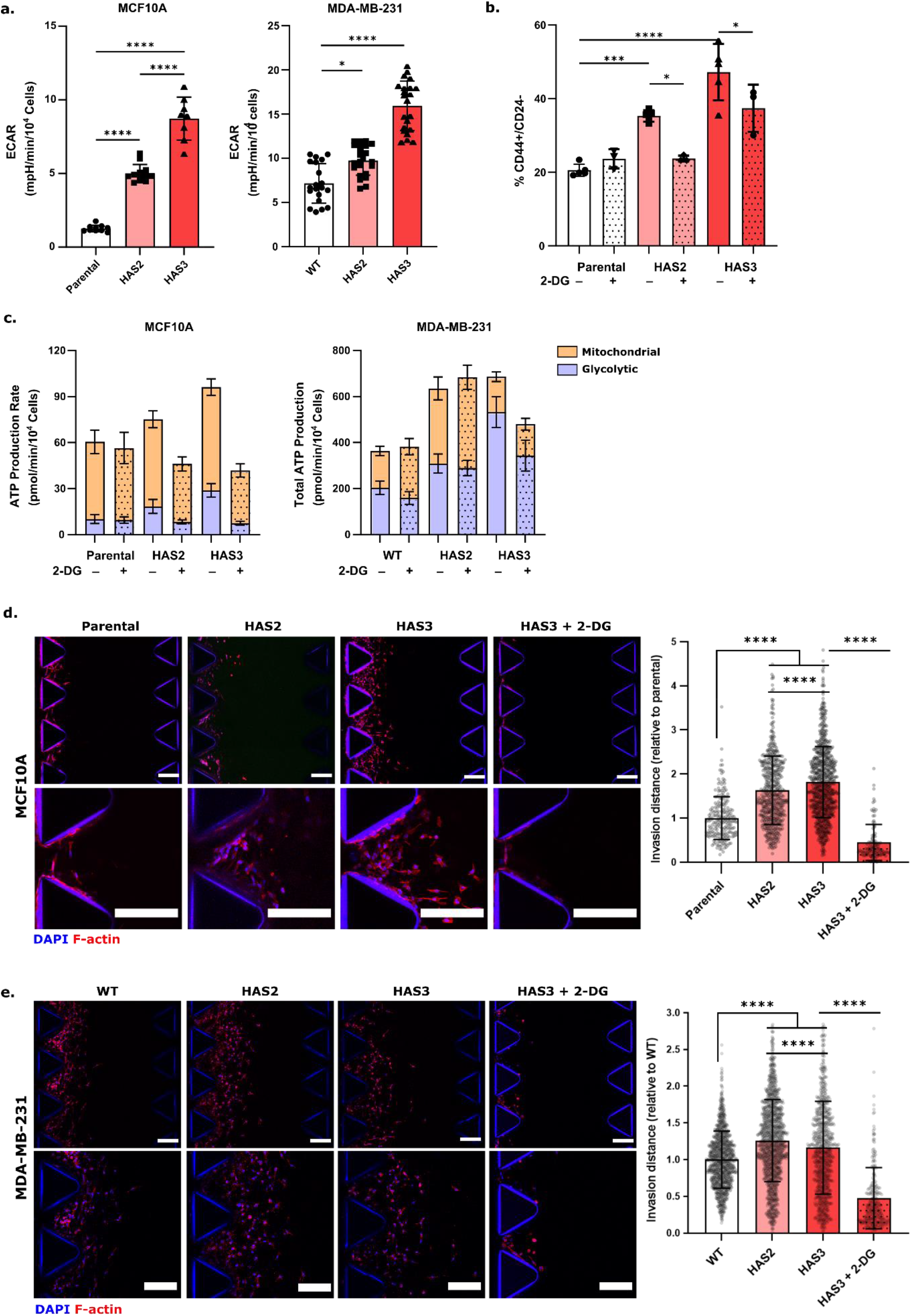
Increased glycolytic metabolism necessary for HA production stimulates stemness and invasion of breast cancer cells. **a)** Extracellular acidification rate measurements of parental MCF10A or HA overproducing cells using the Agilent Seahorse XF Analyzer. **b)** Percentage of CD44^+^/CD24^-^ cells of parental MCF10A or HA overproducing cells with or without 20mM 2-DG as measured by flow cytometry. **c)** ATP production rate of either MCF10A or MDA-MB-231 cells overexpressing HAS2 or HAS3 pre-treated with or without 2-DG (20mM for MCF10A, 25mM for MDA-MB-231) as measured using the Real-Time ATP Rate Assay in the Agilent Seahorse XF Analyzer. (n ≥ 8) **d)** Invasion of the MCF10A and **e)** MDA-MB-231 HA overproducing cells. Representative immunofluorescence images and corresponding quantification of invasion after 5 days are shown (n = 3 devices per condition). Scale bar: 200 μm, * p< 0.05, ** p<0.01, *** p<0.001, **** p<0.0001

### HA Overproduction Transcriptome Changes Predict Worse Patient Survival

As our data implied that HAS2 and 3 overexpression induced more invasive tumor cell phenotypes we next tested if and how these findings may correlate with differences in clinical prognosis. To this end, RNA sequencing was conducted on the MCF10A cells overexpressing HAS2 and HAS3 and their parental control to identify transcriptional changes that would allow us to interrogate the contribution of HA overproduction to stemness, invasion, and patient outcomes. Principle component analysis (PCA) and hierarchical clustering indicated that the transcriptome of HA overproducing cells differed significantly from their parental control (Fig. 6a, b, Supplementary Fig. 6a, Supplementary File 2). Gene set enrichment analysis (GSEA) with the Hallmarks gene sets from the Molecular Signature Database revealed enrichment of a wide range of pathways for both HAS2 and HAS3-overexpressing cells, including those previously associated with stemness and invasion such as NF-κB^42^, hypoxia^8^, PI3K signaling^43^, IL6-STAT3 signaling, and reactive oxygen species^22,23^ (Fig 6c). Since HIF1α signaling has been previously implicated in an HA-dependent increase in the CSC phenotype^17^, we performed GSEA for HIF1α target genes^44–48^. Interestingly, neither HAS2 nor HAS3 transcriptomes were enriched for HIF1α target genes (Fig. 6d), which have previously been suggested as drivers of HA-dependent stemness^17^. These results further support our findings that the cellular phenotypes investigated in this study are due to broader metabolic and energetic changes and cannot be attributed to glycolysis-driven changes in hypoxia-related signaling.

**Figure 6:**
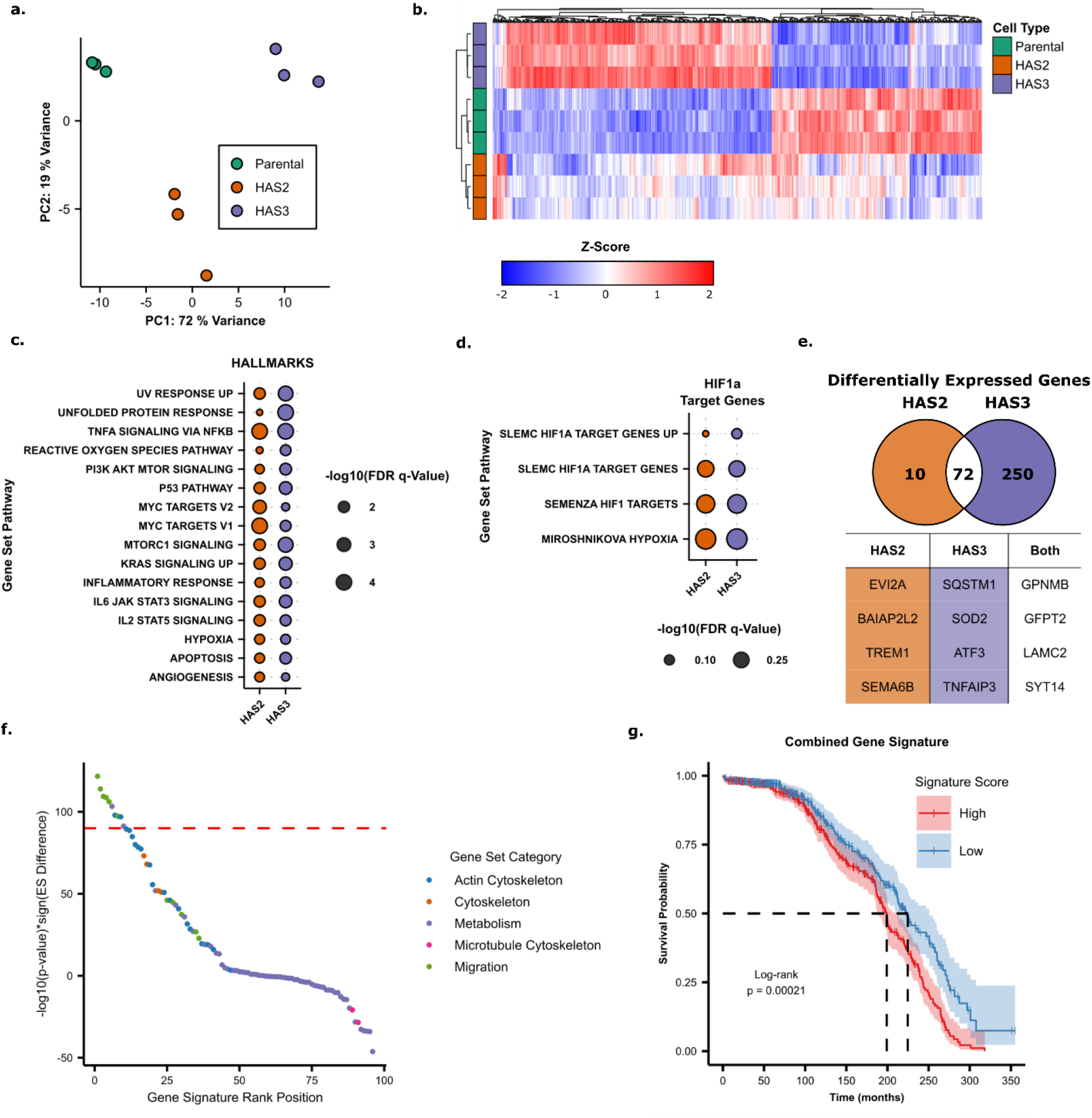
Gene signatures of HA overproducing cells correlate with worse patient prognosis regardless of HAS isoform. **a)** Principle Component Analysis plot of RNA sequencing results of the MCF10A or HA overproducing cells. **b)** Heatmap of z-score normalized gene expression of differentially expressed genes with hierarchical clustering of both samples and genes. **c)** Gene Set Enrichment Analysis of the Hallmarks gene signature database for genes upregulated in either HAS2 or HAS3 overexpressing cells contrasted against Parental MCF10A cells. **d)** Gene Set Enrichment Analysis of published HIF1α target genes for HAS2 or HAS3 overexpressing cells. All results are not significant. (FDR q-values ≥ 0.25) **e)** Venn diagram displaying the total number of significantly increased genes (FDR ≤ 0.05, log2(Fold Change) ≥ 1) in either HAS2 or HAS3 overexpressing cells. The top 4 most significant genes excluding HAS2/3 are shown in the table. **f)** Ranked enrichment analysis of a curated list of migration and metabolism gene sets for tumors from the METABRIC cohort. The degree of enrichment is determined as the -log10(p-value) multiplied by the sign of the difference in enrichment scores (ES) of patients stratified into either low or high enrichment of the HA overproduction gene signature. The dashed red line denotes the empirically determined cutoff for non-random enrichment. **g)** METABRIC patient survival probability predicted by the overlapping 72-gene signature of HA overexpressing cells. Patient signature scores were stratified into quartiles, and a log-rank test was used to determine statistical significance.

To determine the clinical relevance of our findings, a 72-gene signature was generated from the genes upregulated in both HAS2 and HAS3-overexpressing cells (log_2_-fold change > 1, p-value < 0.05) compared to their parental control (Fig. 6e, Supplementary Table 1). Overrepresentation analysis of this gene signature revealed no significant enrichment of genes associated with most metabolic pathways from the KEGG database (Supplementary Fig. 6b). While surprising these results can be explained by the fact that metabolism is significantly regulated by posttranscriptional changes^49,50^. Subsequently, we calculated enrichment scores for this HA overproduction gene signature and a curated list of gene signatures involved in migration, cytoskeleton dynamics, and metabolism (Supplementary File 3, Supplementary Fig. 7a) using single sample GSEA (ssGSEA)^51^ for chemotherapy-naïve patients in the METABRIC cohort. Tumorous tissue from patients enriched for our HA overproduction signature were simultaneously enriched for several migration and actin cytoskeleton gene sets after accounting for random associations suggesting increased tumor invasion in these patients. Metabolic gene sets except for glycosaminoglycan synthesis were not enriched in patients consistent with our findings that metabolic pathways were not overrepresented in our HA production signature nor enriched by both HAS2/3 overexpression in MCF10A cells (Fig. 6f, Supplementary Fig. 6b, Supplementary Fig. 7b,c). Importantly, our HA overproduction signature also predicted worse patient survival consistent with their increased expression of invasion-related gene sets (Fig. 6f,g). Gene signatures specific to upregulation of either HAS2 or HAS3 also predicted worse patient survival that was not seen in the parental control (Supplementary Fig. 6c). Collectively, our results suggest that genes associated with increased HA biosynthesis correlate with an enrichment of migratory genes and predict worse patient survival regardless of HAS2/3 isoform and transcriptional regulation of metabolic gene sets.

## Discussion

Although CSCs have been associated with HA production and altered metabolism, the exact nature of these connections to tumor heterogeneity and consequences on tumor cell invasion remain unclear. Because of the breadth of expertise required to probe each of these aspects, it is infeasible to rely on a single model system or analytical technique to perform a comprehensive investigation of these connections. Furthermore, the systems used must be compatible to enable a holistic evaluation of these aspects. Here, we have used a suite of multidisciplinary approaches that include engineered cell lines, *in vitro* 3D cell culture models, computational metabolic modeling, and genomic tools to uncover the relationship between tumor cell phenotype, HA production, metabolic reprogramming, and 3D invasion. We demonstrate that CSCs can increase glycolytic and oxidative metabolism simultaneously and that the resulting changes in HA production support 3D invasion by supporting more efficient ATP production. Moreover, we identified that these cellular changes correlated with a gene expression signature that was predictive of patient survival.

Our data suggests that cell states associated with increased glycolysis enrich for highly invasive, HA-producing CSCs. Interestingly, HA produced by CSCs is primarily retained on the cell surface rather than excreted into the surrounding environment (Fig. 3, Supplementary Fig. 3). Consequentially, the retention of HA on the cell surface contributes to glycocalyx thickness whose biophysical properties impact the interactions between cells and their surrounding environment including cell-extracellular matrix interactions necessary for migration^18,52^. In particular, HA-dependent changes of the glycocalyx can promote adhesion-independent or ameboid migration by altering the friction required for force generation against extracellular structures^53^, which may further decrease the energy needed for migration. Alternatively, changes in glycocalyx thickness impact surface receptor diffusion patterns and accessibility to impact both adhesion-mediated and receptor tyrosine kinase-mediated signaling^54–56^. Further studies to delineate the contributing biophysical properties are needed to determine the influence of HA on predominant migration mode.

Throughout the metastatic cascade, HA has been implicated in promoting invasive phenotypes and to support survival of circulating cells in the vasculature^19–21^. While HA-dependent changes of cancer malignancy and stemness have been primarily attributed to upregulation of HAS2^15,31^, our findings indicate that HAS3 similarly promotes a stem-like state (Fig. 3). Indeed, our results that 2-DG inhibited the stem-like phenotype in both HAS2/3-overexpressing mammary epithelial cells suggests that the metabolic reprogramming enacted by HA overproduction may be a central regulator of the tumor stem-like phenotype (Fig. 5b). Interestingly, exogenous degradation of HA produced by cancer cells promotes glucose uptake that can further promote migration^56^. This mechanism possibly provides a positive feedback mechanism by which increased HA production regardless of HAS isoform stimulates 3D invasion. Further studies are necessary to decouple the metabolic programming associated with HA production and degradation on the CSC phenotype.

To interrogate alterations in other metabolic pathways associated with glycolysis and oxidative phosphorylation, we utilized FBA to predict changes in metabolic fluxes induced by stemness. FBA is especially proficient in enabling the study of cancer metabolism as it simulates metabolic phenotypes based on real-world constraints such as cell growth rate and glucose uptake^39,57^. The FBA model developed here indicated that stem-like cells exhibited an increased capacity for HA production, and that this increase in HA production was associated with broad metabolic alterations. Interestingly, our model suggested that although the conversion of glucose to pyruvate is decreased in stem-like cells when maximizing HA production, the production of both lactate and acetyl-CoA from pyruvate is increased. An alternative source of pyruvate are the malic enzymes, which convert malate to pyruvate while producing the reducing agent NADPH^58^. Indeed, our FBA model indicated an increase in the flux through malic enzyme 2 (ME2) (Supplementary File 1), which has been previously associated with the loss of cellular senescence and increased tumorigenesis^59,60^. Although any connection with ME2 will need to be experimentally verified in future experiments, the FBA model developed here was able to provide additional insights into the metabolic alterations associated with stemness-related HA production.

The energetic demands associated with HA-mediated invasion can induce broad metabolic changes. For example, increased HA production rapidly depletes UDP-sugar substrates, which cancer cells may compensate for by increasing glycolysis to maintain flux into the HBP and thus, the pool of UDP-sugars^17^. Our FBA model corroborates this phenomenon and provides further insight into metabolic states associated with CSCs such as an increased HA production capacity. The resulting metabolic phenotypes associated with overexpressing HAS2 or HAS3 in cells suggest that the glycolytic demand dominates over possible changes in oxidative metabolism previously associated with stemness^23,61,62^. Furthermore, our finding that HA overproduction induces phenotypic and transcriptomic changes that correlate with invasion may help explain why glycolytic, mesenchymal breast CSCs localize to the leading edge of tumors and worsen patient prognosis^23,36^. Together, these results suggest that HA production enables a more invasive, malignant CSC metabolic phenotype.

Transcriptomic analysis of HAS2 and HAS3 overexpressing MCF10As indicated an enrichment of multiple stemness-associated gene sets (e.g. IL6-JAK-STAT3 signaling and hypoxia), but metabolic genes were not differentially expressed. This discrepancy with our experimental observations may be explained by how metabolic changes are not only regulated transcriptionally but also by enzyme activity levels, localization, and substrate availability^49,50,63,64^. Furthermore, cytoskeletal rearrangement, which is critical for invasion, can independently control glycolytic flux by mediating enzyme degradation^65^ or sequestration^50^. Our results suggest that the glycolytic CSC phenotype associated with HA overproduction is not regulated transcriptionally, but whether this is enacted by cytoskeletal dynamics or changes in the relative activity of different metabolic enzymes requires further investigation.

## Acknowledgements

Thank you to all members of the Fischbach lab for valuable feedback and discussion of the research. The authors would also thank Dr. Andrea Di Micheli, David McKellar, and Dr. Ben Cosgrove for assistance with RNA-sequencing data processing. The work described was supported by National Cancer Institute through the Center on the Physics of Cancer Metabolism (1U54CA210184), the National Science Foundation (NSF) through Graduate Research Fellowship Program awarded to A.A.S and L.M.R. (DGE-1650441), the U.S. Department of Education through the Graduate Assistance in Areas of National Need (GAANN) Fellowship awarded to M.L.T., the Breast Cancer Coalition of Rochester, and the Cornell NanoScale Science & Technology Facility (CNF), a member of the National Nanotechnology Coordinated Infrastructure (NNCI) supported by the National Science Foundation (Grant NNCI-2025233). This work utilized facilities at the Cornell Institute of Biotechnology including the Cornell University Biotechnology Resource Center (BRC) Imaging Facility (RRID:SCR_021741) for image acquisition using the Zeiss LSM 710 Confocal Microscope (NIH 1S10RR025502), the BRC Genomics Facility (RRID:SCR_021727) for sequencing, and the BRC Flow Cytometry Facility (RRID:SCR_021740) for cell sorting with BD Biosciences FACSAria Fusion. Mass spectrometric analysis of targeted metabolomics was performed at the Weill Cornell Medicine Proteomics and Metabolomics Core Facility.

## Author Contributions

A.A.S., M.L.T., and C.F. designed the study. A.A.S. and M.L.T. conducted most of the experiments. M.V. and D.D. conducted extracellular metabolomics for FBA analysis. L.M.R. and M.P. performed FACS for HA production. J.K., L.H., and M.P. generated plasmids for HAS overexpression. L.M.R. and A.A.S. generated cell lines. J.V. and M.L.T. performed FBA analysis. A.A.S., M.L.T., and C.F. analyzed the data and wrote the manuscript. All authors discussed the results and commented on the manuscript.

## Materials and Methods

### Cell Culture

MCF10A (ATCC) cells were cultured in DMEM/F12 supplemented with 5% Horse Serum, 1% penicillin/streptomycin, 10 μg/mL Insulin, 0.5 μg/mL Hydrocortisone, 100 ng/mL Cholera toxin, and 20 ng/mL human epidermal growth factor (EGF). MDA-MB-231 (ATCC) and the GFP-NANOG MDA-MB-231 (a kind gift from Dr. Ofer Reizes^27^) cells were cultured in DMEM supplemented with 10% FBS and 1% penicillin/streptomycin. HAS overexpressing cells were cultured in their respective medias containing 1 μg/mL doxycycline (Santa Cruz Biotechnology). For glycolytic inhibition studies, cells were treated with media containing 2-deoxyglucose (MilliporeSigma) matching the glucose concentration in the media (25mM or 20mM for DMEM, DMEM/F12 respectively) for 24 hours unless otherwise noted.

### Generated Cell Lines

cDNAs for human HAS2 and HAS3 were generated and cloned into the lentiviral vector pLV HygroR tetOn to create stably transduced, tetracycline-inducible MCF10A cell lines as previously described^52^. cDNAs for human HAS2 and HAS3 were also fabricated and inserted into a tetracycline-inducible PiggyBac expression vector to generate pPB huHAS2-IRES2-mScarlet-IRES2-NeoR and pPB huHAS3-IRES2-mScarlet-IRES2-NeoR through custom gene synthesis (Twist Biosience). Generation of the MDA-MB-231 HAS overexpressing cells was conducted using either the pPB huHAS2-IRES2-mScarlet-IRES2-NeoR or pPB huHAS3-IRES2-mScarlet-IRES2-NeoR or with the Piggybac transponase using the Nucleofector Cell Line Kit V (Lonza). Stably transfected cells were selected using 1 μg/mL puromycin (MilliporeSigma) or 800 μg/mL G418 (ThermoFisher). MDA-MB-231 HAS overexpressing cells were then sorted post-selection on mScarlet expression levels.

### FACS and Flow Cytometry

GFP-NANOG MDA-MB-231 cells were trypsinized and resuspended at 10×10^6^ cells/mL in FACS buffer (2.5% FBS/PBS, 2mM EDTA) and processed on the BD FACSAria Fusion Cell Sorter. The top 5% and bottom 5% of the GFP+ fraction were designated as GFP^High^ and GFP^Low^ respectively, while the non-GFP expressing population were designated as GFP^Null^. Cells were sorted into cell culture media, recounted, seeded, and allowed to rest for 24 hours before use in experiments.

Cell sorting for targeted metabolomics was conducted on GFP-NANOG MDA-MB-231 cells stained for HA using Alexfluor-568 (ThermoFisher) conjugated HA binding protein (HABP, 40 μg/mL, MilliporeSigma). The high HA (top 5%) or low HA (bottom 5%) fractions were sorted using a BD FACSAria Fusion Cell Sorter.

ALDH activity was determined for MDA-MB-231 and MCF10A cells using the Aldefluor™ Assay (STEMCELL Technologies) according to manufacturer instructions with incubation conducted for 30min at 37°C. Analysis was conducted on a BD Accuri C6 Plus analyzer.

The fraction of CD44+/CD24− cells was determined by trypsinizing and resuspending cells in FACS buffer at 10×10^6^ cells/mL followed by incubation with antibodies against human CD44 (APC-conjugated, Clone G44-26, 1:5, BD Biosciences) and CD24 (PE-Cy7-conjugated, Clone ML5, 1:20, BD Biosciences). Gates were determined using the isotype controls mouse anti-IgG2b κ (APC-conjugated, Clone 27-35, 1:5, BD Biosciences) and mouse anti-IgG2a κ (PE-Cy7-conjugated, Clone G155-178, 1:20, BD Biosciences). Cells were analyzed on a BD Accuri C6 Plus Analyzer.

### Invasion and Migration Assays

To prepare microfluidic invasion assays, rat tail Type I collagen (Corning) was neutralized with 1 N NaOH and diluted with 1X DMEM to a final concentration of 2.5 mg/mL. Before neutralization and dilution, 10× DMEM was added to collagen as a pH indicator to a final concentration of 10% v/v. Microfluidic 3D cell culture devices (AIM Biotech) were then injected with the 2.5 mg/mL rat tail Type I collagen solution into the center channel of the chip. The chips were then incubated at 4°C for 30 minutes, followed by incubation at 37°C for 30 minutes to complete collagen polymerization. For MDA-MB-231 invasion, DMEM containing 1% FBS and 1% penicillin/streptomycin was injected into the left flanking media channel, and DMEM containing 10% FBS and 1% penicillin/streptomycin was injected into the right flanking media channel. For MCF10A invasion, MCF10A media devoid of Horse Serum and EGF was injected into the left flanking media channel, and fully supplemented MCF10A media was injected into the right flanking media channel. 10 μL of a 2.5×10^5^ cell suspension of GFP-NANOG MDA-MB-231 or MCF10A cells were then injected into both ports of the left channel (20 μL total). Cells were allowed to invade through the hydrogel channel for 3 - 5 days before fixation in a 4% paraformaldehyde solution. Media was exchanged every 24 hours. After fixation, chips were stained, imaged, and individual cell invasion distance was measured using ImageJ by segmenting nuclei.

To assess random migration ability, GFP-NANOG MDA-MB-231 cells were plated on fibronectin-coated (30 μg/mL) glass 96 well plates and placed in an Incucyte S3 (Sartorius) live cell imaging system. Images were obtained in 20-minute intervals over 24 hours. Individual cell tracking was performed using ImageJ to determine migration velocity (motility) and random migration paths.

### Immunofluorescence

GFP-NANOG MDA-MB-231 cells were plated on fibronectin-coated (30 μg/mL) glass coverslips at 2500 cells/cm^2^. Cells were then fixed in 4% paraformaldehyde (PFA)/PBS (w/v) for 20 min at room temperature, blocked with 1% bovine serum albumin (BSA)/PBS (w/v) for 1 hr at room temperature, and incubated overnight with AlexaFluor-568 conjugated hyaluronic acid binding protein (HABP, 13.3 μg/mL, MilliporeSigma) at 4°C. Afterwards cells were incubated with DAPI (2.5 μg/mL, ThermoFisher) for 30 min at room temperature.

For invasion experiments, devices were fixed in 4% paraformaldehyde (PFA)/PBS for 30 min at room temperature, permeabilized with 0.1% Triton X-100/PBS (v/v) for 15 min at room temperature, blocked with 1% bovine serum albumin (BSA)/PBS (w/v) for 1 hr at room temperature, and incubated with DAPI (2.5 μg/mL) and either AlexaFluor-568 (ThermoFisher) or AlexaFluor-647 (ThermoFisher) phalloidin to visualize F-actin.

### Confocal Microscopy and Image Analysis

Images were acquired on a Zeiss LSM 710 confocal microscope with either a LD LCI Plan-Apochromat 25×/0.8 Imm Korr DIC M27 or C-Apochromat W M27 10×/0.45 objective. Images were analyzed using ImageJ with custom scripts. Briefly, for single cell intensity measurements, z-stacks were sum projected and cells were segmented based on HA intensity. Cell clusters were manually corrected to individual cells, while overlapping or edge-located cells were excluded from analysis.

For invasion experiments, the C-Apochromat W M27 10x/0.45 objective was used at 0.6× zoom, and a z-stack was obtained along a 2 mm length in the center of the device. Z-stacks were maximum intensity projected, and invasion was assessed based on nuclei displacement along the x-axis (across the hydrogel region). Invasion distance was normalized to the appropriate control in each set of replicates per experiment, and data was pooled together across device replicates to obtain averages per condition.

### Metabolic Analysis

To obtain an initial measure of glycolytic ability, sorted GFP-NANOG MDA-MB-231 cells were seeded in 24 well plates at 10,000 cells/cm^2^ in 1 mL of media. Glucose concentration was measured using a GlucCell Glucose Monitoring System (CESCO Bioengineering), and lactate concentration was obtained using a colormetric lactate assay kit (Sigma).

Targeted metabolomics was conducted on GFP-NANOG MDA-MB-231 cells 48 hours post sorting for HA production. Media was collected and non-adherent cells were pelleted at 500 × g for 4 minutes. Meanwhile, 0.5 mL of 80% methanol (MetOH) was added onto adherent cells and incubated at −80°C, After aspirating supernatant, the cell pellet was resuspended in 0.5 mL of 80% MetOH and added to the respective well of adherent cells in plate and incubated at −80°C for 15 minutes. Cells were then scraped using a cell scraper and collected into an Eppendorf tube and pelleted at 20,000 × g for 10 minutes at 4°C. Supernatant was then transferred to 2 mL screw-cap vial and dried overnight under vaccum at room temperature. Following overnight drying, samples were then dried for 2.5 hours in SpeedVac SPD 1030 at room temperature and then stored at −80°C until analyzed by Weill Cornell Medicine Proteomics and Metabolomics Core Facility.

Real-time changes in metabolism were tested using a Seahorse XFe96 Analyzer in conjunction with the Seahorse XF Glycolysis Stress Test Assay Kit and the Seahorse XF Real-time ATP Assay Rate Kit (Agilent). Manufacturer instructions were followed to perform each of the assays. Wildtype and modified MDA-MB-231 cells were seeded on a Seahorse XFe96 Cell Culture microplate at 20,000 cells per well in standard media and allowed to attach overnight. For experiments with MCF10A, cells were seeded at 20,000 cells per well. HAS overexpressing cells had media changed 24 hours after seeding to include 1 μg/mL doxycycline. Media was then changed to the specific Seahorse assay media, and plates were prepared according to manufacturer instructions for each assay kit. Relevant metabolic values from each assay were calculated using template worksheets provided by Agilent. After the assays were complete, DNA was extracted from each well using Caron’s Buffer (25 mM Tris-HCl, 0.4 M NaCl, 0.5% (w/v) sodium dodecylsulfate), and total DNA content measured using the fluorometric Quantifluor dsDNA Assay (VWR) and converted to cell number for normalization.

### Flux Balance Analysis (FBA)

To generate a computational model of metabolic fluxes in sorted GFP-NANOG MDA-MB-231 cells, cells were sorted as described above and seeded into 24 well plates at 10,000 cells/cm^2^. Cells were allowed to attach overnight before a fresh media change. Media was collected 72 hours after the initial media change and stored at −80°C before performing extracellular metabolomics to measure levels of glucose, lactate, and the 20 amino acids. Glucose was measured using Contour next EZ Blood Glucose Monitoring System (Ascensia Diabetes Care) using 5 μL as the sample volume. Lactate and amino acid concentrations were assayed using an Acquity UPLC H-Class System equipped with QDa and tunable UV (TUV) detectors controlled by Empower 3 software (Waters Corporation). Specifically, extracellular amino acids were analyzed using a Waters AccQ-Tag Ultra Derivatization Kit (Waters) according to the manufacturer’s recommendations. Derivatized samples were injected onto an AccQ-Tag Ultra C18 column (1.7 μm, 2.1 mm × 100 mm, Waters) and detected by an Acquity TUV detector (Waters) at 260 nm. Amino acids were identified by known retention times of standards and concentrations were determined by comparison with calibration standard curves. For the lactate measurements, samples were first deproteinized by treatment with an equal volume of trichloroacetic acid followed by centrifugation at 12,000 × g for 10 minutes. 200 μL of the supernatant was then combined with 600 μL of acetonitrile (ACN) before injecting 2 μL into the LC-MS system. Lactate was quantified using a standard curve ranging from 0.05mM to 1mM. Separations were performed on an Acquity UPLC BEH Amide Column (1.7 μm, 2.1 mm x 100 mm, Waters). Solvent A consisted of 50:50 ACN:Water and solvent B consisted of 95:5 ACN:Water. The solvent gradient started at 0.01% solvent A and 99.9% solvent B, raised to 40% A in 0.5 minutes, further raised to 70% A in 1.5 minutes, and returned to initial conditions over 0.1 minute and held for 3 minutes to re-equilibrate the column. The flow rate was set to 0.6 mL/min, the autosampler was set to 5°C, and the column was set to 50°C. The mass-to-charge ratio (m/z) of lactate was 88.9. Analysis was performed in negative ion mode with a cone voltage of 15V and probe temperature of 600°C.

Extracellular metabolomics data of all 20 amino acids, lactate, and glucose was then used to constrain a flux balance analysis by imposing bounds on allowable fluxes. Growth rate and O2 fluxes were constrained from cell counts and Seahorse data respectively. The model used in this study was implemented in the Julia programming language^66^, where the linear programming problem was solved using the GNU Linear Programming Kit (GLPK) package (https://www.gnu.org/software/glpk/). The stoichiometric matrix and metabolic growth requirements were derived from a previously developed Core Cancer model^41^. The objective function was set to maximize HA production subject to experimentally estimated rates of uptake, secretion, and cell growth to define the phenotype of interest; 1000 simulations were performed for each condition. A subset of the FBA results representing the energetic pathways of interest was incorporated into a graphical representation (Fig. 4). The full FBA results can be found in Supplementary File 1.

### Limited Dilution Assay

Cells were serially diluted and seeded into ultra-low attachment 96-well plates (Corning) in 200 μL of serum-free DMEM/F-12 containing 2% B27, 10 ng/mL basic fibroblast growth factor, 20 ng/mL epidermal growth factor, 10 μg/mL insulin, and 1 μg/mL doxycycline. Sphere formation was assessed after two weeks of culture, and the number of spheres was counted using a phase-contrast microscope. The stem cell frequency was determined using the extreme limited dilution algorithm^67^.

### RNA -Sequencing

Parental (NeoR-rtTA) and HA overproducing MCF10A cells were seeded on 10-cm dishes at 5000 cells/cm^2^ and allowed to adhere overnight. Media was refreshed to include 1 μg/mL of doxycycline to induce HAS2 and HAS3 expression and cultured for an additional 48 hours. RNA was isolated using the Qiagen RNeasy kit according to manufacturer instructions. RNA libraries were prepared using the Illumina TruSeq RNA Kit, and single-ended 75bp read lengths were sequenced on the Illumina NextSeq 500 system.

### Sequence Alignment and Gene Set Enrichment Analysis

Reads were trimmed using TrimGalore version 0.4.4 (https://www.bioinformatics.babraham.ac.uk/projects/trim_galore/) and aligned to the human reference genome GRCh38 (ENSEMBL) using STAR version 2.6.0a^68^. Reads of genomic features were counted using featureCounts^69^, and differential gene expression was determined using DESeq2^70^. Differentially expressed genes in HA overproducing cells were defined as a log-2 fold change greater than 1 and an adjusted p-value less than 0.0001 compared to the parental cells. Genes common to both HAS2 and HAS3 were combined to generate the HA overproducing gene signature for survival analysis. Individual HAS2 and HAS3 gene signatures were defined as the differentially expressed genes with log 2-fold change greater than 2 or 5, respectively, and a p-value less than 0.0001.

Gene set enrichment analysis (GSEA) was conducted with a ranked list generated by taking the sign of the fold change multiplied by the log-10 of the adjusted p-value. The list was inputted to the GSEA Java applet (http://software.broadinstitute.org/gsea/index.jsp) using the GSEAPreRanked tool and the Hallmarks gene sets from MSigDB v.7.0. Gene sets were considered significantly enriched with a p-value and FDR value ≤ 0.05.

### Patient Survival and Enrichment Analysis

Patient data from the METABRIC cohort was extracted from the Cancer Genomics Data Server. Patients were limited to those having not received chemotherapy. Gene signature scores were calculated using the single-sample GSEA (ssGSEA)^71^ with the GSVA package^72^. The top and bottom quartiles of the ssGSEA scores for the 72-gene HA overproduction signature were designated as high and low scores, respectively, for survival and enrichment analysis. Kaplan-Meier survival analysis was conducted with the *survival* package in R using a Cox proportional hazard model with statistical significance determined using a log-rank test. Stratified patients were further analyzed for enrichment of curated lists of migration, cytoskeletal, and metabolic gene signatures obtained from the MSigDB v7.0. To correct for random associations, 300 randomly selected gene signatures with the same number of genes (15-500 genes) as gene sets in the curated list had ssGSEA enrichment scores additionally calculated. An empricial cumulative distribution function was established using the *ecdf* function in R and a p-value cutoff was determined where 95% of values fell below.

### Statistical analysis

All experiments were performed with at least three independent biological replicates unless otherwise noted. Pairwise comparisons were conducted using a Mann-Whitney U test unless otherwise noted. Multiple comparisons were evaluated with either a Kruskal-Wallis Test or two-way ANOVA with Dunn’s post hoc analysis. Results were considered statistically significant with a p-value less than 0.05. Unless otherwise noted, all data points are plotted mean +/− the standard deviation. All statistically analysis was performed using GraphPad Prism v9.3 or R.

