## Supplementary Figures for "Convergent approaches to delineate the metabolic regulation of tumor invasion by hyaluronic acid biosynthesis"

|  |  |  |  |  |
| --- | --- | --- | --- | --- |
| GRAMD1B | PCSK5 | SAA2 | SYT14 | PTAFR |
| SH2D2A | JAG1 | LRP4 | S100A8 | MTSS1 |
| TLL1 | RHOV | DSG3 | IQGAP2 | CHRNA9 |
| LAMA3 | TNNT1 | FHOD3 | NCAM1 | B3GNT3 |
| LAMC2 | PTN | SLC37A2 | BTBD11 | SATB1 |
| SLC2A3 | PTPRZ1 | ADAM19 | WNT7A | AMTN |
| EYA2 | MDFI | BICDL1 | XDH | ARL4C |
| COL17A1 | HES1 | GPNMB | LAD1 | HEPACAM2 |
| TP63 | IL1A | CLCA2 | S100P | CES1 |
| CA12 | ITGB6 | GPAT3 | STEAP1 | CES1P1 |
| FYB1 | GPR68 | AMIGO2 | GJB2 | RPSAP52 |
| CD82 | BMP2 | RDH16 | RNASE7 | MMP28 |
| FAT2 | GFPT2 | CDH11 | SCG5 |  |
| HSD17B2 | KANK4 | COL6A2 | FAM83B |  |
| HEPH | PTPN22 | SLC47A1 | BDKRB2 |  |

**Supplementary Table 1: Hyaluronic Acid Overproduction Gene Signature.** The 72-gene signature of upregulated genes associated with HA overproduction found in both HAS2 and HAS3 overexpressing cells.

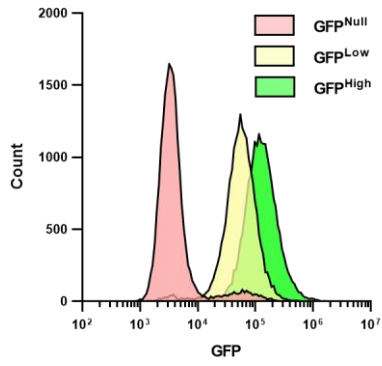

**Supplementary Figure 1: GFP fluorescence of sorted GFP-NANOG MDA-MB-231 as measured by flow cytometry after 5 days of culture.**

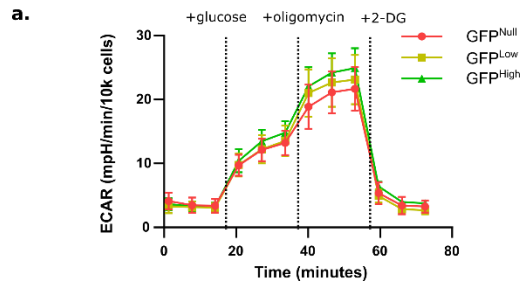

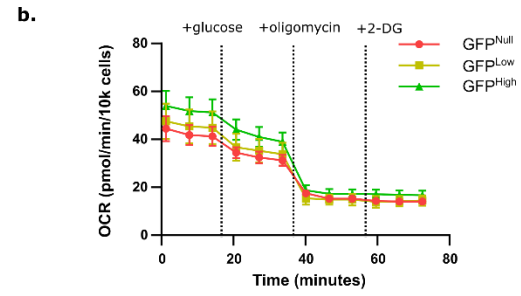

**Supplementary Figure 2: Real-time traces of the metabolic state of sorted GFP-NANOG cells. a) Extracellular acidification rate (ECAR) b) Oxygen consumption rate (OCR) of sorted cells undergoing the Glycolysis Stress Test normalized by cell number over time.**

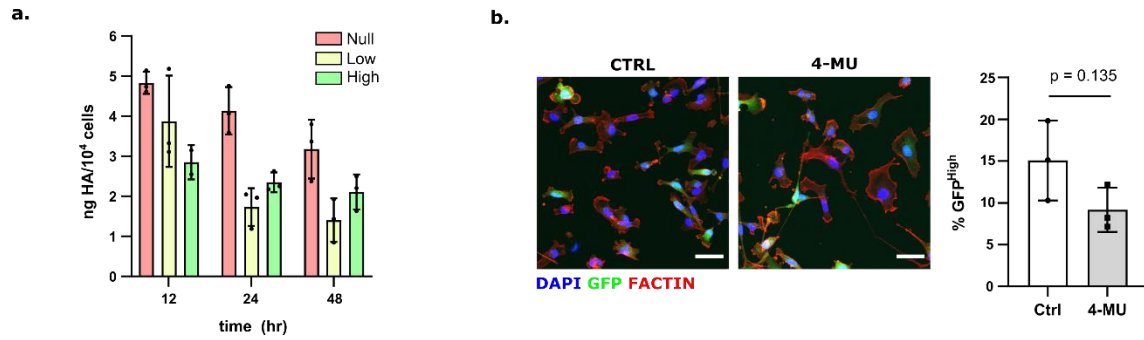

**Supplementary Figure 3: HA Production of GFP-NANOG cells. a)** HA ELISA of sorted GFP-NANOG MDA-MB-231 (n = 3). **b)** GFP high fraction of cells treated with or without 4-MU as quantified by image analysis (n = 3). Scale bar = 50 μm.

a.

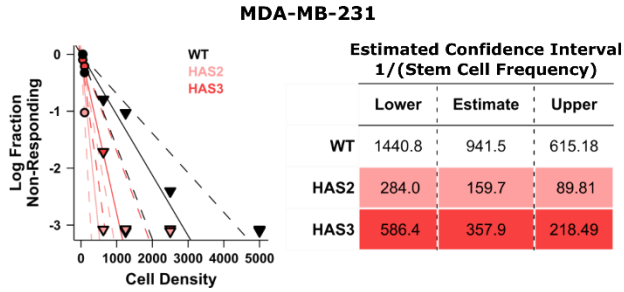

b.

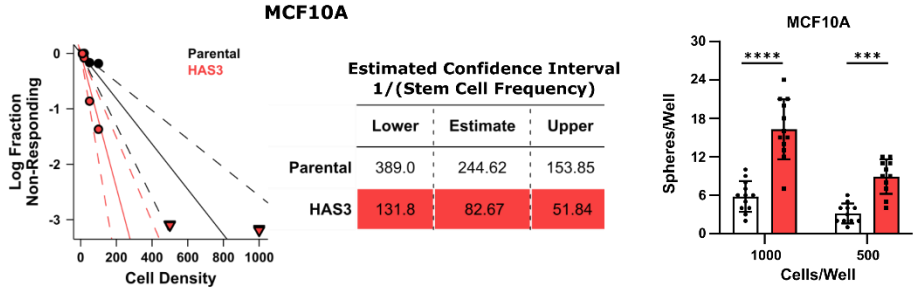

c.

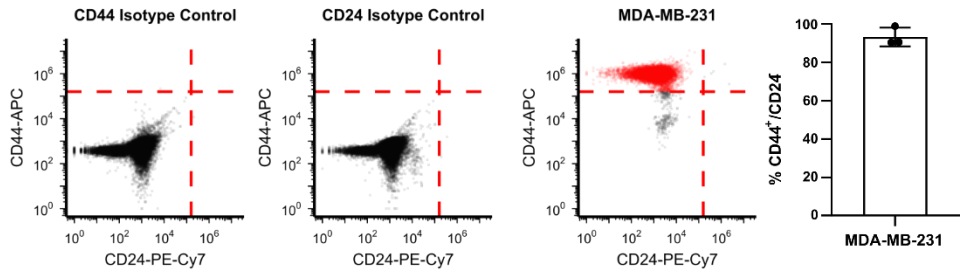

d.

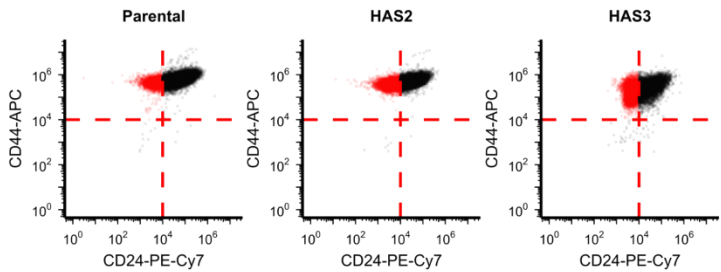

e.

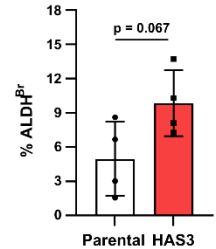

### Supplementary Figure 4: Identification of CSC enrichment induced by HA

overproduction. a) Limited dilution assay results of WT MDA-MB-231 or HAS2/3

overexpressing cells (n = 11). Left, Plots showing the fraction of wells without sphere formation against the cell seeding density. The slope of the solid line represents the estimated stem cell frequency, and dashed lines the 95% confidence interval as determined by the extreme limited dilution analysis algorithm. Downward pointed triangles represent densities with a zero negative response. Right, Table containing the estimated and 95% confidence interval of the stem cell frequency. b) Limited dilution assay for MCF10A parental and HAS3 overexpressing cells (n = 11). Right, Quantification of the number of spheres in wells seeded with 500-1000 cells. c) Flow cytometry of CD44/CD24 stained MDA-MB-231 cells with isotype controls. Average measurements of the CD44<sup>+</sup>/CD24<sup>-</sup> population in MDA-MB231 (n = 3) to the right. d) Flow cytometry plots for the CD44/CD24 in parental or HA overproducing MCF10A cells. Dashed red lines represent gating thresholds based on isotype controls, and red areas signify the CD44<sup>+</sup>/CD24<sup>-</sup> population. e) Flow cytometry analysis of the aldehyde dehydrogenase bright fraction (ALDH<sup>Br</sup>) of HAS3 MCF10A as measured by the Aldefluor assay (n = 4). \*\*\* p<0.001, \*\*\*\* p< 0.0001

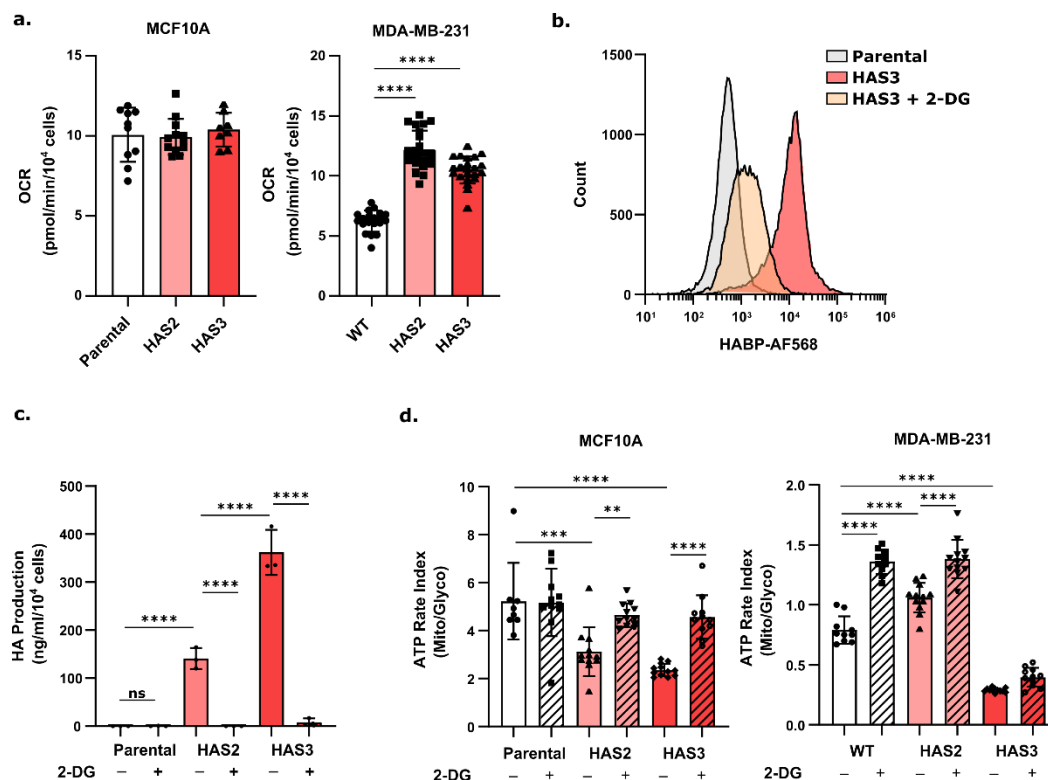

**Supplementary Figure 5: Increases in the CSC phenotype due to increased HA biosynthesis are tied to glycolysis.** **a)** Oxygen consumption rate of HAS overexpressing MCF10A and MDA-MB-231 as measured using the Glycolysis Stress Test Assay in the Agilent Seahorse XF Analyzer. **b)** Flow cytometry analysis of cell-surface associated HA of either parental or HAS3 overexpressing MCF10A cells with or without 20mM of 2-DG using fluorescently tagged HA binding protein (HABP-AF568). **c)** Measurements of HA secretion into cell culture media of HA overproducing MCF10A cells treated with or without 20mM 2-DG as measured with an HA ELISA normalized by cell number (n = 3). **d)** ATP rate index (Mitochondrial ATP/Glycolytic ATP) of HAS overexpressing MCF10A and MDA-MB-231 with or without 2-DG (20mM for MCF10A, 25mM for MDA-MB-231) as measured using the Real-Time ATP Rate Assay in an Agilent Seahorse XF Analyzer (n ≥ 8). \*\* p<0.01, \*\*\* p<0.001, \*\*\*\* p< 0.0001

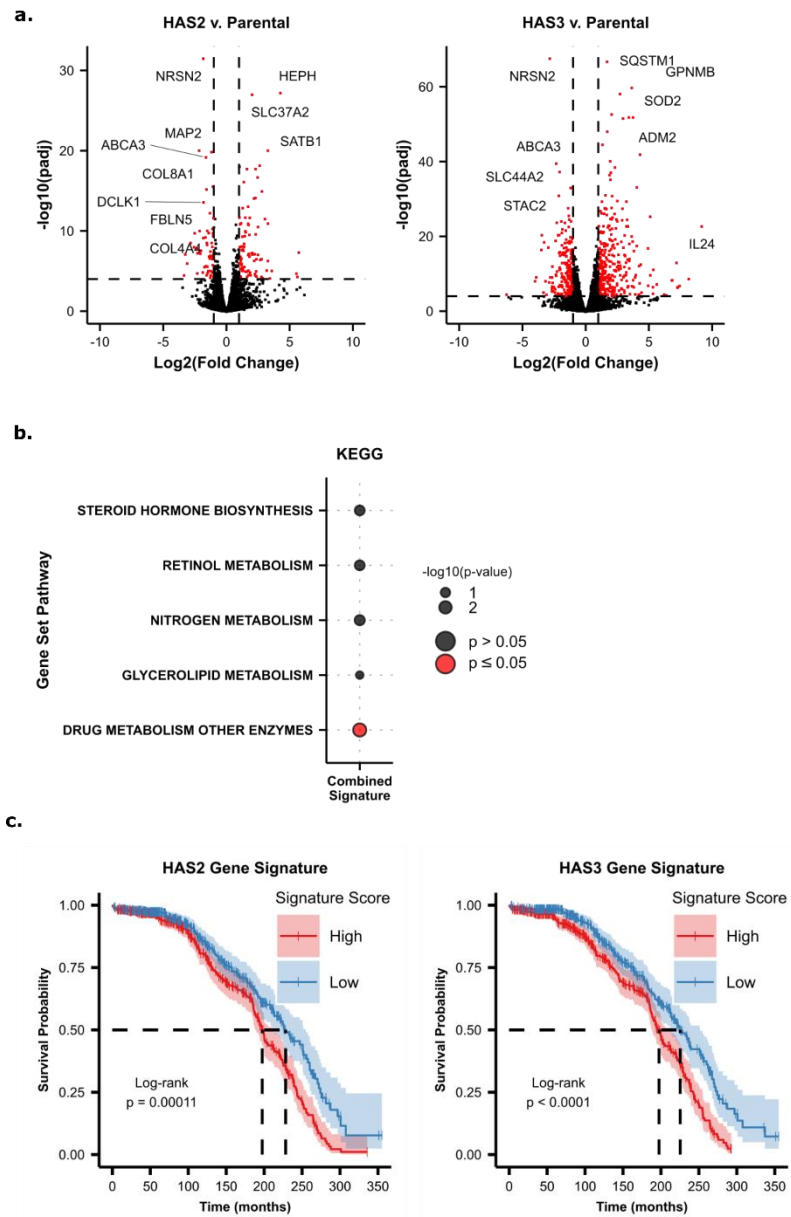

**Supplementary Figure 6: Expanded RNA-sequencing results.** **a)** Volcano plots depicting gene expression differences between either the HAS2 or HAS3 cells with their parental controls. Red dots indicate differentially expressed genes with a $\log_2$ fold change $\geq 1$ and p-adjusted value $\leq 0.05$. **b)** Over-representation analysis of metabolic gene signatures from the KEGG database of the 72-gene signature of HA overproducing cells. **c)** Overall survival analysis of chemotherapy-naïve patients in the METABRIC cohort stratified into the upper and lower quartiles enriched for either the HAS2 or HAS3 upregulated genes.

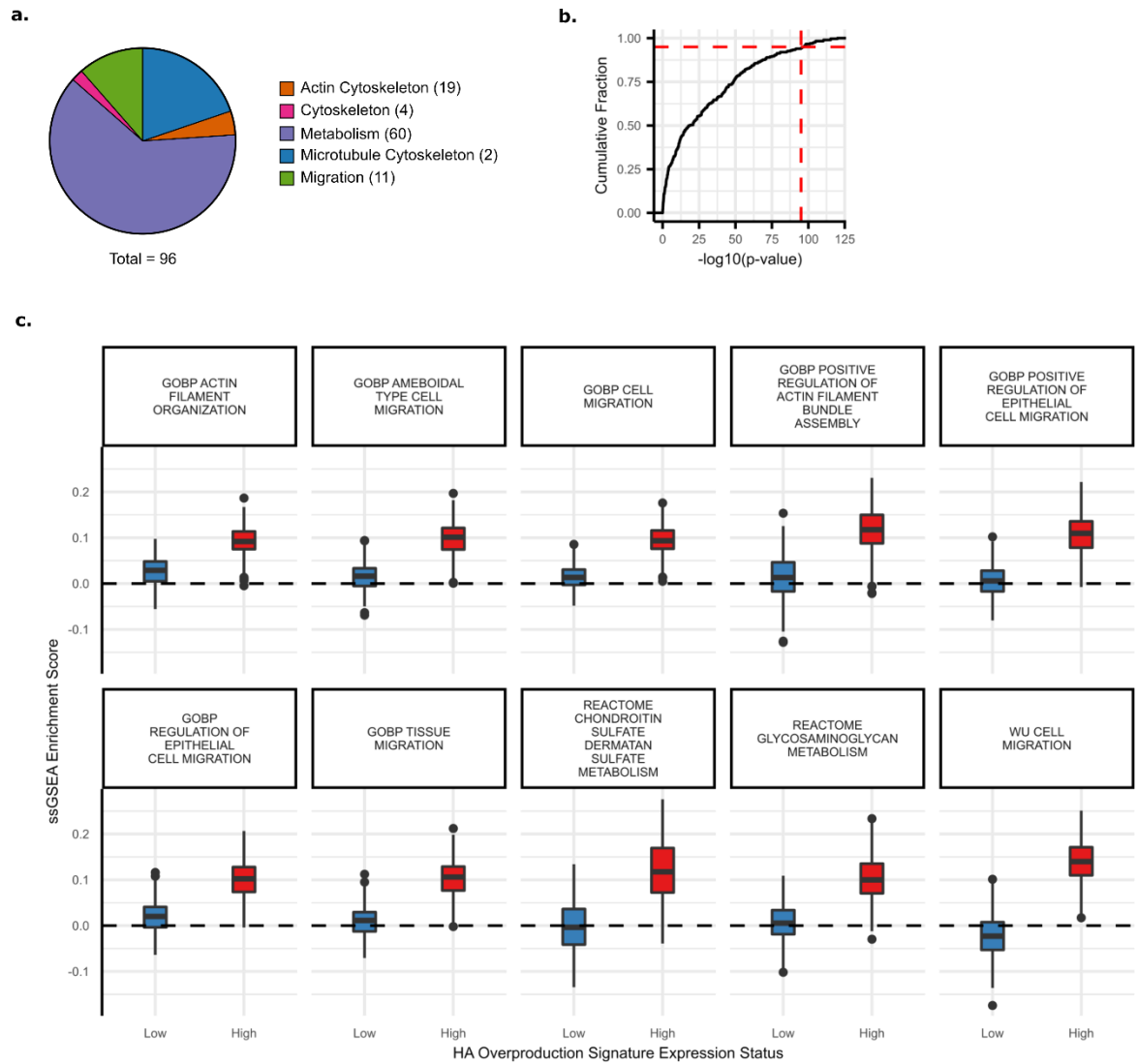

**Supplementary Figure 7: High HA overproduction gene signature enrichment correlates with migration gene sets in the METABRIC cohort.** **a)** Pie graph showing the distribution of gene set categories in the curated gene list. Number of gene sets in each category is indicated in legend. **b)** The empirical cumulative distribution function of p-values for differences in the enrichment score of 300 randomly selected gene sets from the Molecular Signature Database. Patients were stratified based on their enrichment status of the HA overproduction signature. Dashed lines signify the cutoffs for strong enrichment not due to random association ($p\text{-value} \leq 1e-90$). **c)** Single sample GSEA enrichment scores for the top 10 curated migration and metabolism associated gene sets that were concurrently enriched in patients with a high HA overproduction enrichment.
